# Modulating SPARC Expression in Mesenchymal Stem Cells Improves Secretome-Mediated Skin Regeneration and Wound Repair

**DOI:** 10.64898/2026.03.07.710278

**Authors:** A. Lombardi, J. Smucler, M.B. Palma, A. Iribarne, A. La Greca, M.N. García, G.E. Sevlever, S.G. Miriuka, C.D. Luzzani

## Abstract

Mesenchymal stem cells (MSCs) have garnered significant attention over the past three decades due to their robust regenerative potential, primarily mediated by their paracrine activity by releasing soluble bioactive factors and extracellular vesicles (EVs). The MSC secretome plays a pivotal role in wound healing by influencing cellular migration, inflammation, angiogenesis, extracellular matrix (ECM) remodeling, and re-epithelialization. SPARC (Secreted Protein Acidic and Rich in Cysteine), a multifunctional ECM glycoprotein involved in tissue repair and remodeling, regulates key processes such as cell migration, proliferation, angiogenesis, and survival. Despite its known role in ECM dynamics, the impact of SPARC expression on the regenerative properties of MSCs remains underexplored.

In this study, we hypothesized that SPARC overexpression in MSCs enhances their secretome’s regenerative capacity. Using lentiviral systems, we generated SPARC-overexpressing (+SPARC) and SPARC-knockdown (KD-SPARC) MSCs to investigate SPARC’s role in wound healing. Conditioned media (CM) derived from these MSCs were analyzed *in vitro* for their effects on human skin keratinocytes and fibroblasts. Our results revealed that SPARC expression significantly influences cell-specific migration and cell cycle. Furthermore, in an *in vivo* wound healing model, CM from +SPARC MSCs accelerated regeneration, while SPARC absence in MSCs CM delayed the healing process.

These findings underscore the critical role of SPARC in modulating MSC secretome composition and enhancing its regenerative efficacy. This study highlights SPARC as a promising therapeutic target for the development of advanced regenerative therapies aimed at improving cutaneous wound healing outcomes.

## INTRODUCTION

Mesenchymal stem cells (MSCs) are multipotent stromal cells with remarkable regenerative potential, making them a cornerstone of regenerative medicine research over the past three decades. Their therapeutic effects are largely mediated by their secretome, a complex mixture of soluble factors, cytokines, and extracellular vesicles (EVs) that influence various cellular processes. This paracrine activity of MSCs has been shown to play a critical role in promoting tissue repair by modulating inflammation, stimulating angiogenesis, enhancing cellular migration and proliferation, and facilitating extracellular matrix (ECM) remodeling^1–3^. These properties make MSCs a promising candidate for developing therapies targeting cutaneous wound healing, involving the intricate interplay of inflammation, angiogenesis, tissue formation, re-epithelialization, and ECM remodeling^3–5^.

Wound healing is a dynamic process that often requires therapeutic intervention to achieve optimal outcomes, particularly in chronic wounds or severe injuries^6^. MSCs have demonstrated the ability to enhance the healing process by promoting a regenerative microenvironment^7–9^. Furthermore, advances in bioengineering have focused on modifying MSCs to improve their regenerative capabilities. Such modifications aim to optimize the composition and potency of their secretome, a cell-free therapeutic approach that holds great promise for overcoming the challenges associated with cell-based therapies, including immunogenicity and scalability^10^.

A protein of particular interest in this context is SPARC (Secreted Protein Acidic and Rich in Cysteine), also known as BM-40 or osteonectin. SPARC is a multifunctional matricellular (extracellular matrix-associated) glycoprotein^11^. Unlike its name (osteonectin) might suggest, SPARC expression is not limited to bones, it is also present in diverse tissues including non mineralized tissues, in platelets and in muscles. Such wide distribution correlates with its roles during embryogenesis as well as during tissue repair, cell turnover, cellular differentiation and remodeling, which are key steps in tissue regeneration. Interestingly, the situations in which SPARC is overexpressed are mainly those requiring regeneration, either to repair tissues (injury) or adapt to tissue changes (obesity, exercised muscle, etc.)^12,13^. Furthermore, the regeneration process requires the implication of numerous cellular organelles and the use of energy. In this context, SPARC has been shown to be implicated in a variety of metabolic functions, such as glucose tolerance improvement, while it is also required for both glucose homeostasis maintenance and insulin secretion^12^. In addition, SPARC has interesting roles within the inflammatory processes as it has anti-inflammatory properties, and modulates factors related to the immune response^12,14^.

All these highlighted properties point to SPARC as a regeneration factor. It not only has significant roles in tissue repair or development but contributes directly and indirectly to generating a “positive” biological environment that optimizes regeneration. Despite its known regenerative functions, the specific impact of SPARC on the secretome of MSCs and how this could contribute to MSCs intrinsic regenerative properties has not yet been described.

This study aims to investigate the effects of SPARC overexpression and knockdown in MSCs and their regenerative capacity. By analyzing the paracrine effects of SPARC-modified MSCs on human skin keratinocytes and fibroblasts *in vitro*, as well as in an *in vivo* mouse wound healing model, we seek to elucidate SPARC’s role in enhancing MSC-mediated wound regeneration. Understanding how SPARC influences MSC functionality could pave the way for developing more efficient, targeted therapies for wound healing and tissue repair.

## RESULTS

### Modulation of SPARC Expression in WJ-MSCs

To modulate SPARC expression in WJ-MSCs, we utilized a lentiviral system due to its high transduction efficiency and ability to maintain cell viability. For SPARC overexpression, the SPARC mRNA sequence from WJ-MSCs was cloned into the pLJM1-CMV-Puro lentiviral vector. To achieve SPARC knockdown, a specific short hairpin RNA (shRNA) sequence was cloned into the pLKO.1-Puro vector (Fig. 1a), followed by transduction into WJ-MSCs. After transduction and antibiotic selection, no significant increase in cell death was observed, and cell morphology remained consistent with non-transduced controls, indicating high cell viability and absence of cellular stress (data not shown).

**Figure 1.**
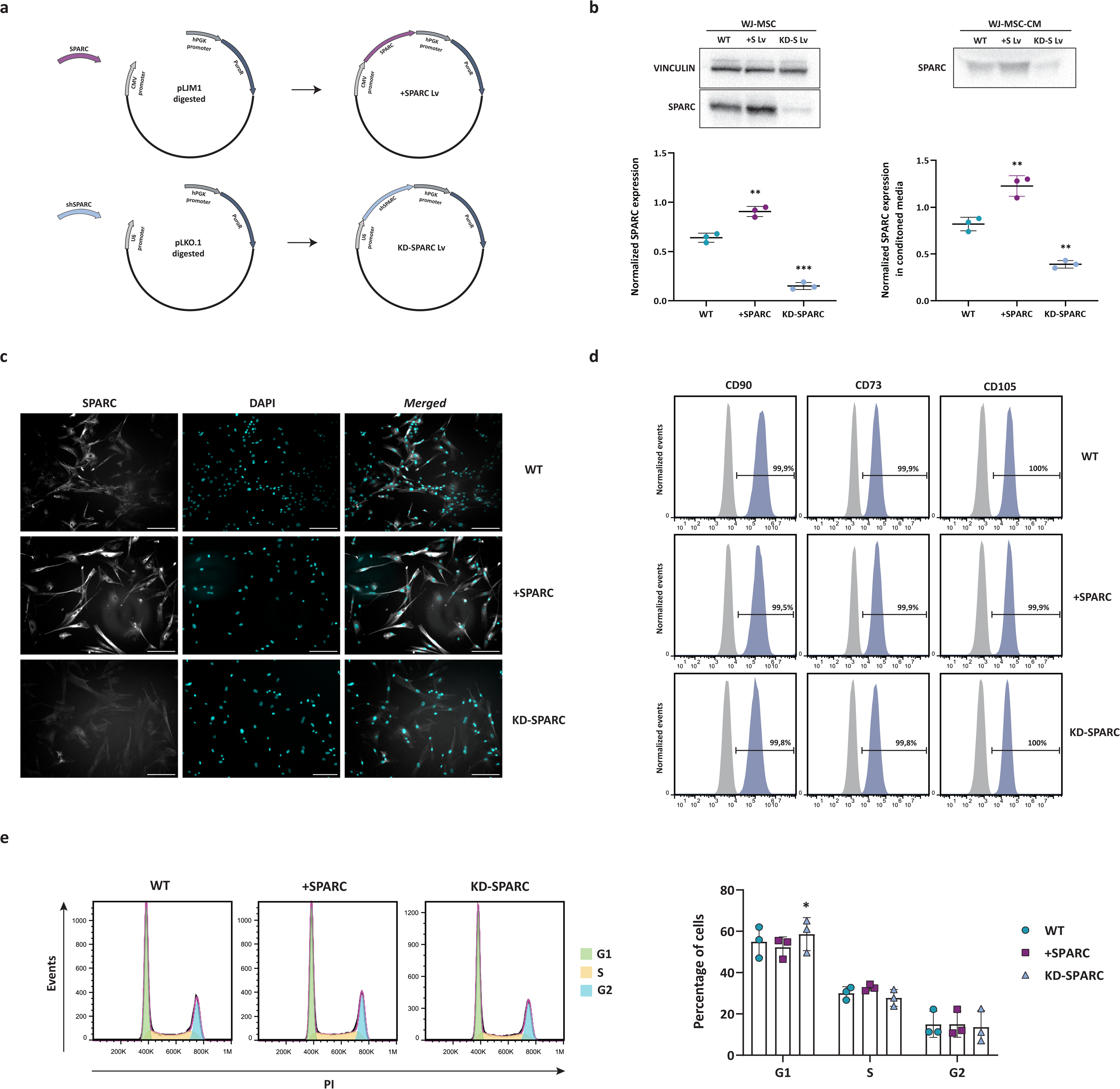
Generation and Characterization of SPARC-Modified WJ-MSCs. a: Schematic graph showing the inserts and backbone plasmids used to build the +SPARC Lv (upper graph) and KD-SPARC Lv (lower graph) constructs **b:** SPARC expression levels in WJ-MSC lysates (left panel and graphic) and their conditioned media (CM) (right panel and graphic) assayed by SDS-PAGE and Western blot. Cell lysates and CM were obtained from wild-type (WT) WJ-MSCs or those transduced with +SPARC Lv or KD-SPARC Lv. Quantification represents the mean of three independent assays. The blot shown is composed of bands cropped from different parts of the same gel (whole blot is shown in Supplementary Fig.1). **c:** SPARC localization in WT, +SPARC and KD-SPARC WJ-MSC, assayed by immunofluorescence and fluorescence microscopy. Representative images of three different experiments are shown. Scale bar: 200 µm. **d:** MSC identity markers CD90, CD73 and CD105, measured by flow cytometry. Representative plots of three independent experiments. **e:** Cell cycle analysis by propidium iodide incorporation assayed by flow cytometry of WJ-MSC WT, +SPARC and KD-SPARC. Right panel: quantification of three independent experiments on the percentage of cells in each phase. The percentages were calculated with the Un-variate platform of the FlowJo v10.0 software, statistical analysis were made with Graphpad 8.0 software, p<0.05 (*).

Western blot analysis confirmed both successful overexpression and knockdown of SPARC protein (Fig. 1b, Supplementary Fig. 1), with corresponding changes in the levels of secreted SPARC (Fig. 1b, right panel and graph). Immunofluorescence analysis further validated the differences in SPARC expression and showed no changes in its intracellular localization among wild-type (WT), SPARC-overexpressing (+SPARC), and SPARC-knockdown (KD-SPARC) cells (Fig. 1c).

Cellular characterization revealed that all cell groups maintained typical MSC surface markers (CD90, CD73, and CD105) without differences between WT, +SPARC, and KD-SPARC cells (Fig. 1d). The expression of typical negative surface markers remained unchanged (Supplementary Fig. 2), confirming that SPARC modulation did not alter WJ-MSC identity.

Given that SPARC has been implicated in regulating processes such as cell proliferation, we investigated the cell cycle dynamics of the modified cells, as this is a critical aspect of proliferation control. Cell cycle analysis, conducted using propidium iodide (PI) staining and flow cytometry, revealed that all groups exhibited a typical MSC cell cycle distribution, with the majority of cells in the G1 phase and fewer cells in the S and G2 phases (Fig. 1e, left panels). Notably, KD-SPARC cells showed a significantly higher proportion of cells in the G1 phase compared to WT cells, suggesting a reduced proliferation rate. In contrast, no significant differences were observed in the cell cycle profiles of +SPARC cells relative to WT cells (Fig. 1e, right graph).

Furthermore, we evaluated the expression of specific membrane integrins known to be regulated by SPARC in other cell types. These integrins, which are expressed in MSCs, play key roles in signaling pathways that govern cell adhesion and migration. Our analysis showed no significant differences in integrin expression between WT cells and MSCs with modified SPARC expression (Fig. 2).

**Figure 2.**
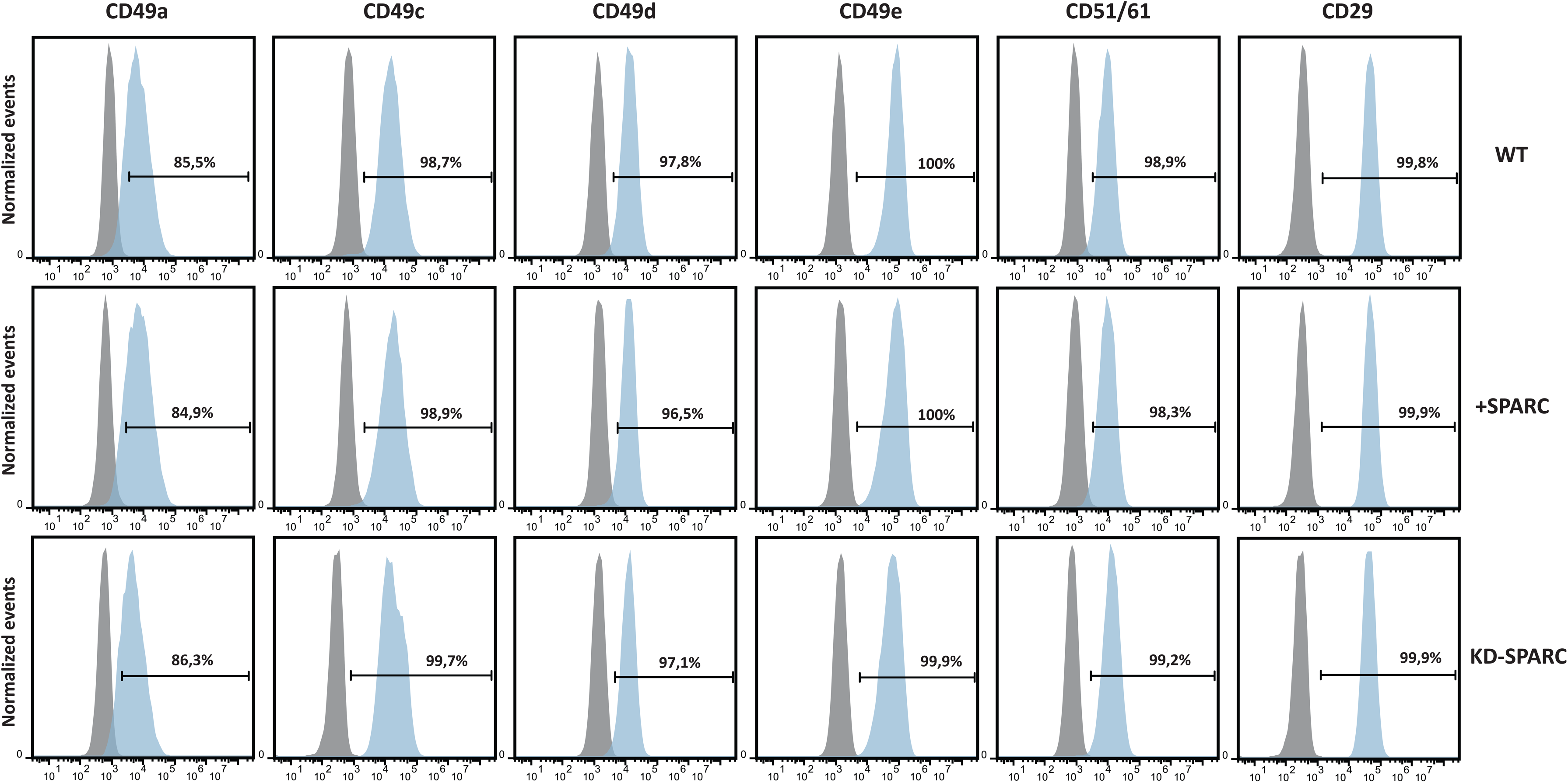
Analysis of Integrin Expression Profile of SPARC-modified WJ-MSCs. Expression of membrane integrins CD49a, CD49c, CD49d, CD49e, CD51/61 and CD29 analyzed by flow cytometry in WT, +SPARC and KD-SPARC WJ-MSC. Plots are representative of three biological replicates. Percentages were calculated with the FlowJo v10.0 software. Statistical analysis was made with Graphpad 8.0 software.

Taken together, these results suggest that SPARC modulation in WJ-MSCs was successfully achieved without compromising cell viability, morphology, or MSC identity. While SPARC knockdown showed a slightly altered cell cycle by an increasing proportion of cells in the G1 phase, overexpression had no significant impact. Additionally, neither up nor downregulation of SPARC expression affected the expression of membrane integrins involved in cell adhesion and migration.

### WJ-MSC +SPARC and KD-SPARC conditioned media induce functional changes in human skin fibroblasts and keratinocytes *in vitro*

MSC secretome, composed of both soluble factors and extracellular vesicles (EVs), represents the primary mediator of MSC-driven biological functions^2^. Its regenerative potential has been demonstrated in various models such as treatment of lung, cerebral, myocardial and hepatic injury, osteoarthritis, corneal and skin wound healing, fibrosis and several other pathologies^2,10^. To explore the influence of SPARC expression on the composition and biological effects of the MSC secretome we collected conditioned media (CM) from cells with SPARC overexpression (+SPARC-CM) and SPARC knockdown (KD-SPARC-CM) and tested the impact of these secretomes on human skin keratinocytes and fibroblasts evaluated through various functional assays.

First, we examined the effect of CM derived from WT, +SPARC, and KD-SPARC WJ-MSCs in an *in vitro* wound-healing model using the immortalized human skin keratinocyte line HaCaT. Confluent keratinocytes were cultured to form a monolayer, and a wound was created on the surface. CM from either WJ-MSCs WT, +SPARC, or KD-SPARC was then added, with the base medium used for MSC conditioning serving as the control. Cells were maintained for 14 hours under standard conditions, with images captured every 30 minutes (Supplementary videos 1, 2, 3 and 4). Using the data acquired from the cell monolayer area on each side of the wound, the percentage of wound area covered was determined as a function of time as shown in Fig. 3a.

**Figure 3.**
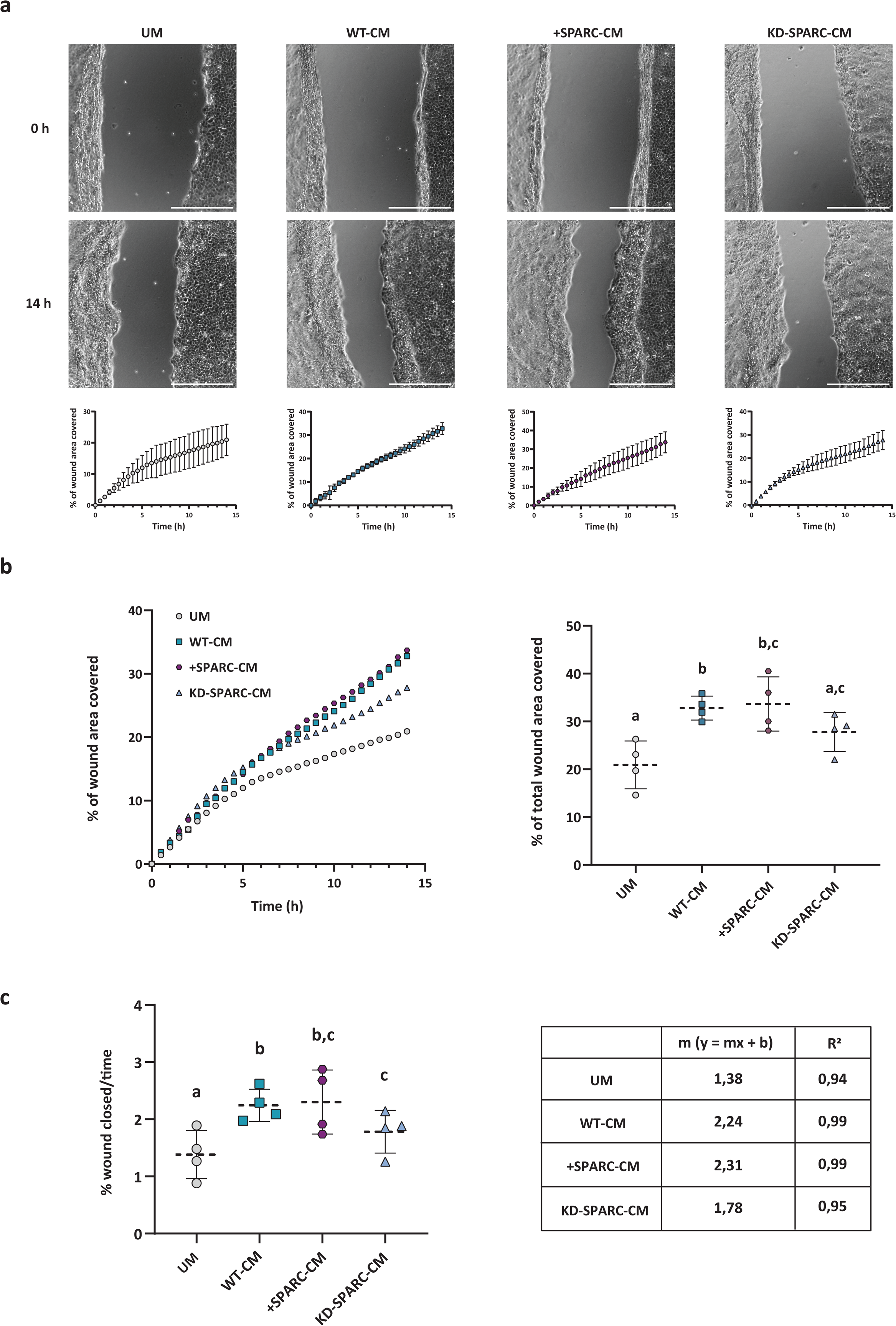
HaCaT migration varies after exposure to WT, +SPARC or KD-SPARC WJ-MSC conditioned media. a: Representative images of wound healing assay. Time 0 and 14 hours after wound performed with HaCaT cells cultivated in conditioned media (CM) of WT, +SPARC and KD-SPARC WJ-MSC. DMEM + 10% FBS, or unconditioned media (UM), was used as a control. Lower graphs show the percentage of wound area covered by the migrating cells on each time point for each condition. Mean values and standard errors are shown. **b:** Mean values of percentage of wound area covered on each time point for all conditions (left panel). Percentage of total wound area covered (right panel). **c:** Percentage of wound closed over time for all conditions (left panel). Slope and R2 values for the linear regression of each curve (right table). For every condition four biological replicates were made. Values and statistical analysis were made with Graphpad 8.0 software. Significant differences are shown as letters, p<0.05.

A detailed analysis of the wound closure velocity curves reveals a slight decrease in the slopes for the unconditioned media (UM) and KD-SPARC-CM conditions, indicating a delay or reduction in cell migration. This trend is not observed when comparing the conditions with WT-CM and +SPARC-CM, where a more linear and consistent migration is evident throughout the evaluated period (Fig. 3a, lower graphs). Figure 3b compares all the curves on the same graph, showing differences in the final percentage of wound area covered.

Quantification of the total average wound area covered at the endpoint shows that keratinocytes migrate to cover a significantly higher percentage of the wound when cultured with WJ-MSC-derived media, regardless of SPARC expression, compared to unconditioned media. When comparing among MSC-derived media, a tendency for greater closure was observed with the WJ-MSC +SPARC medium relative to the others, although it was not statistically significant. Importantly, significant differences were found between the conditions with *wild type* and knockdown conditioned medium, with the latter showing a notably lower percentage of wound area covered (Fig. 3b, right panel), suggesting that the presence of SPARC is important for wound closure.

Finally, we calculated the slope of each curve, representing the average migration velocity, through linear regression analysis. Consistent with previous findings, we observed significant differences in keratinocyte migration velocity in UC compared to mesenchymal-derived media. No significant differences were detected between the WT-CM condition, where the migration velocity of HaCaT cells covered 2.24% of the wound area every 30 minutes, and the +SPARC-CM condition, where the velocity covered 2.30% every 30 minutes. However, in the KD-SPARC-CM condition, the migration velocity was significantly lower than WT-CM, with a wound area coverage of 1.78% every 30 minutes (Fig. 3c). This represents a 20% reduction in migration velocity compared to the other conditions.

Based on the results, it can be concluded that keratinocytes exhibit a pro-migratory effect when exposed to media conditioned by mesenchymal cells, and this effect varies depending on SPARC expression. However, wound healing requires not only cell migration toward the injury site but also their proliferation. To address this, we analyzed the effect of CM on the cell cycle of HaCaT keratinocytes and human dermal fibroblasts (CCD-32Sk) treated with CM derived from WT, +SPARC, and KD-SPARC cells using propidium iodide staining followed by flow cytometry analysis.

The cell cycle profiles of HaCaT keratinocytes were found to be very similar across all conditions. When comparing the proportion of cells in each phase of the cycle, a trend was observed toward a higher percentage of cells in the G1 phase when cultured with CM compared to DMEM. We found that this difference was significantly higher in the KD-SPARC-CM condition. Additionally, differences in the percentages of cells in the S and G2 phases were observed when comparing UM with KD-SPARC-CM (Fig. 4a).

**Figure 4.**
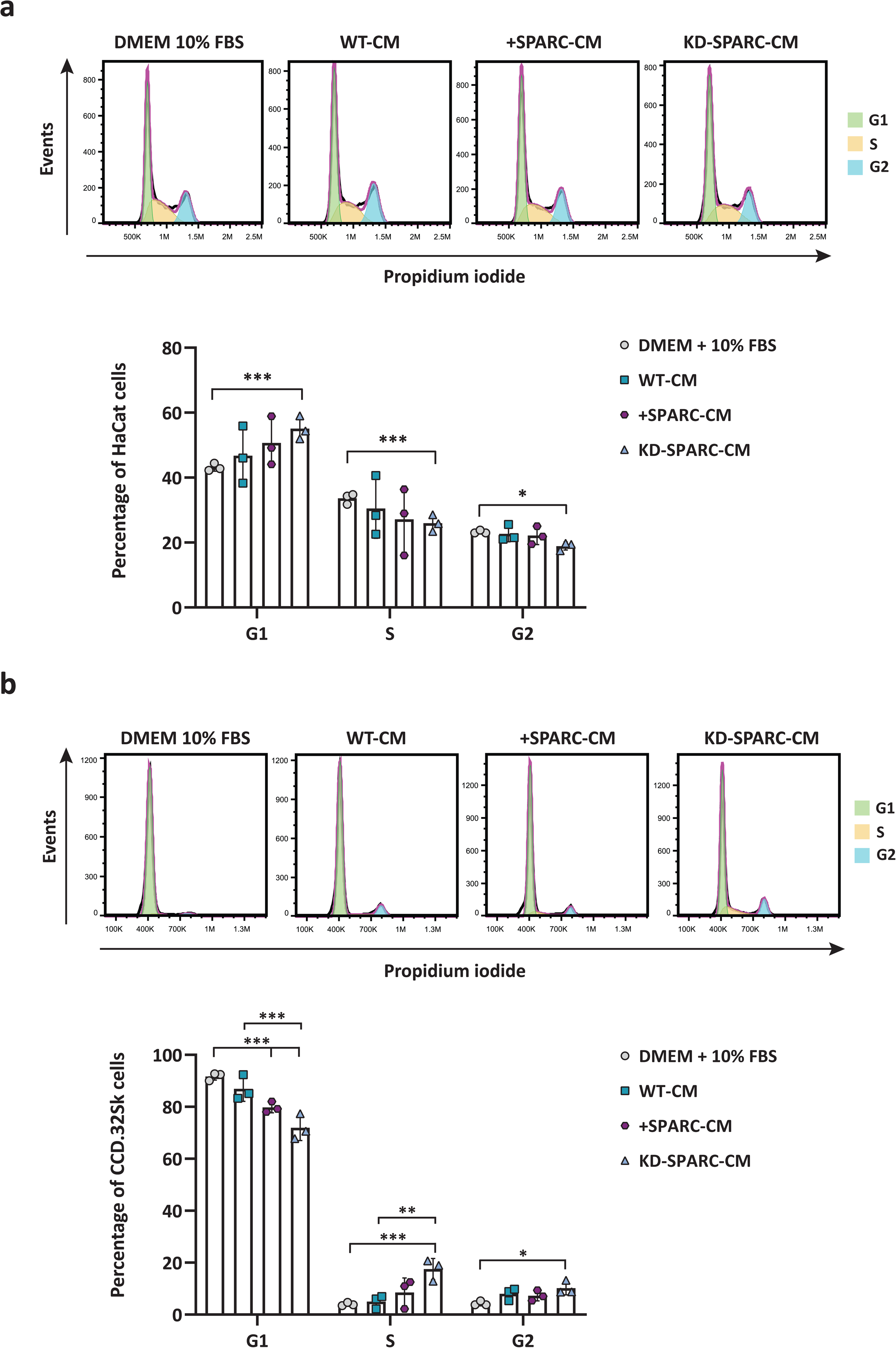
HaCaT and CCD-32Sk proliferation varies upon exposure to WT, +SPARC or KD-SPARC WJ-MSC conditioned media. a: Cell cycle analysis by propidium iodide incorporation assayed by flow cytometry of HaCat cells incubated 24 hours with WT-CM, +SPARC-CM or KD-SPARC-CM. Quantification of the percentage of cells in each phase was performed with the Un-variate platform of FlowJo v10.0 software, statistical analysis was performed with Graphpad 8.0 software, p<0.05 (*), p<0.001 (***). **b:** Cell cycle analysis by propidium iodide incorporation assayed by flow cytometry of CCD-32Sk cells incubated 24 hours with WT-CM, +SPARC-CM or KD-SPARC-CM. Quantification of the percentage of cells in each phase was performed with the Un-variate platform of FlowJo v10.0 software, statistical analysis was performed with Graphpad 8.0 software, p<0.05 (*), p<0.005 (**), p<0.001 (***).

In contrast, fibroblasts exhibited the inverse effect in cell cycle profiles between treatments. We observed a lower percentage of cells in the G1 phase for CM treatments compared to DMEM, with this reduction being significant for +SPARC-CM and KD-SPARC-CM conditions. A similar reduction was found when comparing WT-CM with KD-SPARC-CM. Fibroblasts treated with KD-SPARC-CM displayed a higher percentage of cells in the S phase compared to those cultured in DMEM and WT-CM. Finally, fibroblasts showed an increased proportion of cells in the G2 phase when cultured with KD-SPARC-CM (Fig. 4b).

Altogether, these results suggest that the medium derived from WJ-MSCs also influences keratinocytes and fibroblasts cell cycle, with the effect being more pronounced when the medium is derived from mesenchymal cells with SPARC knockdown. Interestingly, the +SPARC-CM condition produced intermediate values between the WT-CM and KD-SPARC-CM conditions, contrary to the expected outcome. Additionally, it is noteworthy that the observed effects on keratinocytes and fibroblasts are opposite, indicating that the impact of the WJ-MSC secretome may be cell type-specific.

### SPARC Modulation Induces Changes in Gene Expression in WJ-MSCs

Given the functional effects observed when removing SPARC from WJ-MSC’s CM, we sought to evaluate how the modification of SPARC expression in these cells alter their gene expression profile. RNA sequencing (RNA-Seq) analysis revealed significant transcriptomic alterations between wild-type (WT), SPARC-overexpressing (+SPARC), and SPARC-knockdown (KD-SPARC) WJ-MSCs.

Differential expression analyses identified 29 differentially expressed genes (DEGs) in +SPARC versus WT cells and 123 DEGs in KD-SPARC versus WT cells (Fig. 5a, Supplementary Tables 1 and 2). When running a hierarchical clustering algorithm of all 152 DEGs, we found a clustering tendency of biological replicates according to experimental conditions (WT, +SPARC, and KD-SPARC), confirming that SPARC expression significantly shapes the transcriptional landscape of WJ-MSCs (Fig. 5b). Notably, four distinct clusters can be distinguished, reflecting the presence of condition-specific gene expression patterns, as some genes are exclusively expressed in one condition or another. At the same time, certain genes are shared across multiple conditions, indicating overlapping regulatory networks and suggesting both unique and common pathways influenced by SPARC expression.

**Figure 5.**
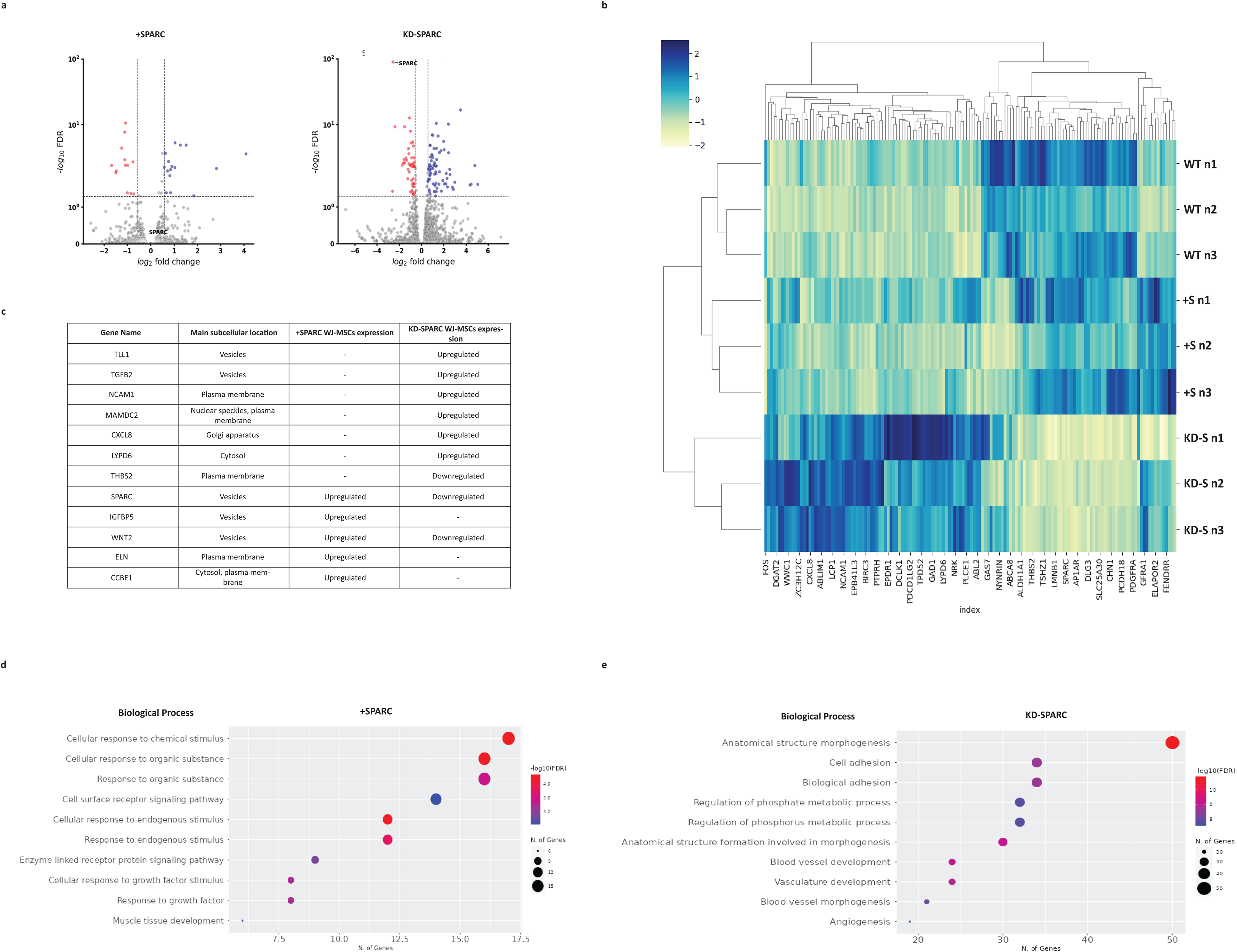
Transcriptomic Impact of SPARC Modulation in WJ-MSCs. a: Volcano plots for RNASeq analysis. 29 differentially expressed genes for WT vs +SPARC WJ-MSC (left panel) and 123 differentially expressed genes for WT vs KD-SPARC WJ-MSC (right panel). Gates at p-value<0.05 and fold-change>1.5. **b:** z-score heatmap for all 152 differentially expressed genes. y-axis shows biological replica for each condition WT, +S (+SPARC) and KD-S (KD-SPARC). **c:** Table: Differentially Expressed Genes Predicted to Encode Secreted Proteins (Identified via Human Protein Atlas Analysis). **d:** Biological Process (BP) gene ontology (GO) enrichment analysis was performed on differentially expressed genes between the WT vs +SPARC condition using ShinyGO. **e:** Biological Process (BP) gene ontology (GO) enrichment analysis was performed on differentially expressed genes between the WT vs KD-SPARC condition using ShinyGO.

We further analyzed the DEGs using the Human Protein Atlas database. This analysis revealed that 12 DEGs encode proteins with secretion potential into the extracellular environment (Fig. 5c). Notably, several of these genes are well-established mediators of intercellular signaling and tissue repair. TGFB2 and CXCL8, both upregulated in KD-SPARC cells, are key regulators of inflammation and angiogenesis, respectively. ELN and IGFBP5, which were upregulated in the +SPARC condition, are associated with extracellular matrix structure and growth factor binding, respectively. Interestingly, WNT2 was the only gene upregulated in the +SPARC condition and downregulated in the KD-SPARC condition. This protein plays a crucial role in Wnt signaling, a pathway involved in cell fate determination, tissue regeneration, and wound healing. Its bidirectional regulation further supports the idea that SPARC expression influences distinct gene networks.

Gene Ontology (GO) Enrichment Analysis using ShinyGO provided additional insights into the biological processes regulated by SPARC-modulated gene expression (Fig. 5c, d). In +SPARC cells, enrichment was observed in processes associated with cellular responses to growth factors. Conversely, in KD-SPARC cells, pathways related to cell adhesion and angiogenesis were notably enriched. These findings demonstrate that overexpressing or knocking down SPARC leads to distinct transcriptional outcomes, underscoring its multifaceted role in modulating cellular functions and the extracellular environment, and suggesting that SPARC is a key player in wound healing-related processes.

So far, we observed that SPARC modulation significantly alters the gene expression profile of WJ-MSCs, influencing key biological processes such as cell migration, proliferation, adhesion, and angiogenesis. These transcriptomic changes likely underline the functional effects observed in fibroblasts and keratinocytes, highlighting SPARC’s pivotal role in regulating the secretome and cellular behavior of WJ-MSCs.

### Conditioned medium from SPARC-Overexpressing or Knockdown WJ-MSCs modulate skin wound healing in Mice

To further examine the impact of SPARC expression on the regenerative capacity of mesenchymal cells, we aimed to evaluate the effect of the secretome from WT, +SPARC, and KD-SPARC mesenchymal cells in wound healing experiments using an *in vivo* model. Immunocompetent C3HS strain mice were used, in which a dermal wound was created using a dermatological punch. After 7 days, a second wound, known as a second-intention injury, was generated in the same area and conditioned medium derived from WT, +SPARC, and KD-SPARC cells was immediately administered via intradermal injections. The wound healing process in the mice was recorded and analyzed up to 21 days after the first injection (Fig. 6a).

**Figure 6.**
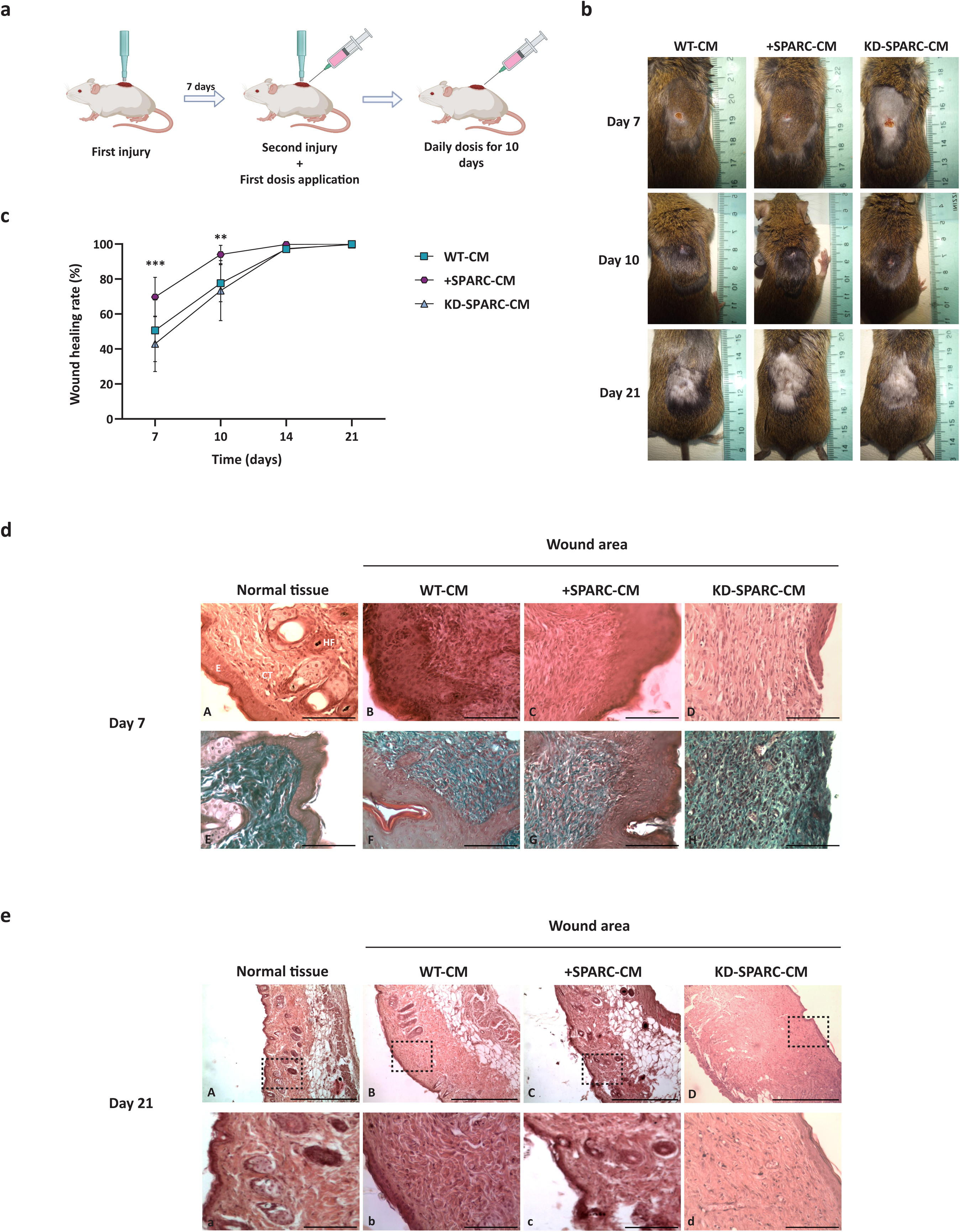
WJ-MSC conditioned media affects mice wound healing in a SPARC dependent manner. a: Scheme of experimental design for the second intention injury test and intradermal injection of conditioned media of WJ-MSC WT, +SPARC and KD-SPARC in immuno-competent C3HS mice. **b:** Representative images of mice at days 7, 10 and 21 after second injury and first CM dosis application. On day 21, mice were shaved to appreciate the injury area. **c:** Quantification of the wound healing rate. The difference between the initial wound area and the wound area at each time point was calculated. The average values are expressed in percentages and the standard errors are shown. Three biological replicates were performed with at least 3 mice per condition. Statistical analysis was made using Graphpad 8.0 software. Significant differences were found at day 7 between WT-CM and +SPARC-CM treatments with a p<0.01 (**); and between +SPARC-CM and KD-SPARC-CM with p<0.001 (***). On day 10 we found significant differences between +SPARC-CM and KD-SPARC-CM, p<0.01 (**). The graph shows the asterisks indicating the largest differences found at that time point. **d:** Representative histological images of samples from mice injected with WT-CM, +SPARC-CM or KD-SPARC-CM taken at day 7 after the first injection. Panel A: normal tissue stained with hematoxylin and eosin (E: epithelium (epidermis), CT: connective tissue, HF: hair follicle); B,C and D: samples of the wound area from mice treated with WT-CM, +SPARC-CM and KD-SPARC-CM stained with hematoxylin and eosin, scale bar: 100 μm. Panels E, F, G and H: representative images of Masson’s trichrome staining of samples from normal tissue and mice injected with WT-CM, +SPARC-CM or KD-SPARC-CM taken on day 7 post-injury, scale bar: 100 μm. **e:** Representative images of hematoxylin and eosin staining of histological samples from mice injected with MC WT, MC +SPARC or MC KD-SPARC taken at day 21 after the first injection. Panels A, B, C and D: normal tissue and mice injected with WT-CM, +SPARC-CM or KD-SPARC-CM respectively, scale bar: 500 μm. Panels a, b, c and d: magnifications of the areas indicated with black quadrants in panels A, B, C and D respectively, scale bar: 100 μm.

Figure 6b shows representative images of animals treated with CM from WT, +SPARC, or KD-SPARC WJ-MSCs on days 7, 10, and 21. The most pronounced macroscopic differences between groups were observed on days 7 and 10. By day 7, mice treated with +SPARC-CM exhibited notably smaller wounds with more advanced epithelialization compared to the other groups. Additionally, these mice showed signs of crust formation over the wounds and increased hair regrowth in the area surrounding the lesions. In contrast, the WT-CM and KD-SPARC-CM-treated groups displayed slower healing, characterized by larger wounds, incomplete epithelialization, and limited hair growth in some animals. An additional control assay was conducted in parallel using unconditioned medium (DMEM + 10% FBS). This group exhibited a noticeably slower wound healing rate compared to those treated with MSCs-derived conditioned media (data not shown).

By day 10, wounds in most +SPARC-CM-treated mice were nearly indistinguishable to the naked eye, with many animals achieving complete wound closure. In comparison, wounds in the KD-SPARC-CM-treated group showed noticeable progress but remained visible. The WT-CM-treated group displayed more advanced healing than the KD-SPARC-CM group but lagged behind the +SPARC-CM group, as wounds in this condition were not yet completely closed. By day 21, nearly all mice from all treatment groups exhibited complete wound closure (Fig. 6b).

To quantify these observations, the wound area was measured on days 7, 10, 14, and 21, and the wound closure percentages were calculated (Fig. 6c). On day 7, the +SPARC-CM-treated group showed a significantly higher healing rate compared to the WT-CM and KD-SPARC-CM groups. Although no statistically significant differences were observed between the WT-CM and KD-SPARC-CM conditions, the latter group exhibited a trend toward a slower healing rate. By day 10, the healing rate in the +SPARC-CM-treated group approached 100% and remained significantly higher than in the KD-SPARC-CM-treated group. From day 10 onward, wound healing progressed rapidly in both the WT-CM and KD-SPARC-CM groups, with most individuals showing almost complete wound closure by day 14. By day 21, no significant differences in wound closure were observed across the groups, as healing was largely complete for all conditions.

To further investigate the macroscopic differences observed among the treatment groups, we analyzed the histological characteristics of the wound-healing process. Skin samples from the lesion area were collected at 7- and 21-days post-treatment, fixed in 10% formalin, processed using standard histological techniques, and stained with Hematoxylin-Eosin.

Histological sections from wounds treated with intradermal injections of WT-CM on day 7 revealed key features of the normal healing process. Regeneration of the epithelial tissue was evident, with an increased number of cell layers compared to normal skin, along with the presence of keratin in the apical region. Beneath the epithelium, the dermis displayed highly cellular connective tissue (CT), characterized by numerous fibroblasts with tapered nuclei, blood capillaries, and abundant collagen fibers (Fig. 6d, panel B). Histological samples taken from the group of mice treated with +SPARC-CM at 7 days post-injury showed no major differences compared to the WT-CM group, although slightly higher cellularity was observed in the superficial dermis, near the epithelium (Fig. 6d, panel C). Additionally, in some cases, the epithelial tissue appeared to have fewer cell layers. Histological analysis of the KD-SPARC-CM-treated group revealed significant differences compared to the WT-CM and +SPARC-CM groups. A marked cellular infiltration was observed in all cases, characterized by an abundance of lymphocytes, identifiable by their small, rounded, pyknotic nuclei. The dermis also showed a high density of blood capillaries and small-caliber vessels. Some individuals exhibited a notable delay in the healing process, with only scar tissue visible at the wound site, exposed to the exterior without epithelial coverage, and an exacerbated inflammatory response (Fig. 6d, panel D).

We aimed to gain deeper insights into the defining features of the emerging connective tissue (CT). Specifically, we focused on the type, arrangement, and bundling of collagen fibers, as these features provide crucial information about the progression and quality of wound closure. To achieve this, we utilized Masson’s trichrome staining, which differentially stains collagen fibers with light green dye. Figure 6d (panels B, C, and D) shows histological sections from wounds treated with WT-CM, +SPARC-CM, and KD-SPARC-CM, respectively, at 7 days post-injury—the time point where the greatest macroscopic differences were observed.

In wounds treated with WT-CM or +SPARC-CM, collagen fibers appeared intermediate in thickness, moderately irregular in arrangement and interspersed among CT cells, resulting in a slightly lax CT. These features indicate a more advanced and organized wound healing process (Fig. 6d, panels F and G). In contrast, wounds treated with KD-SPARC-CM exhibited abundant fibrotic tissue, characterized by collagen fibers that were thinner and arranged more parallel relative to the basement membrane of the epithelial tissue. The fibers also formed thinner bundles compared to the thick fiber bundles typically observed in normal dermal CT (Fig. 6d, panels E and H).

We then analyzed the samples taken at 21 days post-injury, where the wound area had significantly reduced in size in all groups. In mice treated with WT-CM, the epithelium showed partial normalization, with fewer cell layers compared to day 7, although it remained thicker than uninjured skin in a small region of the wound. The underlying CT exhibited abundant collagen fibers that had undergone reorganization, forming thicker bundles arranged in multiple directions, consistent with the irregular dense CT of the dermis. Notably, no regeneration of hair follicles was observed at this time point (Fig. 6e, panel B). As for the +SPARC-CM group, complete skin regeneration was generally observed, including the formation of hair follicles (Fig. 6d, panel C). Most samples in this group showed no trace of injury, and the tissue closely resembled normal skin. Finally, the KD-SPARC-CM treated mice showed that the inflammatory process had been resolved, with no infiltrating cells present. The epithelial tissue resembled that of normal skin, but the underlying CT layer appeared thicker than in the other groups (Fig. 6e, panel D).

These observations indicate that the absence of SPARC delays the regeneration process, as wounds treated with KD-SPARC-CM exhibited a greater proportion of fibrotic tissue and a slower wound healing rate, reflecting impaired and less effective repair. In contrast, wounds treated with +SPARC-CM demonstrated reduced fibrosis, more advanced connective tissue remodeling, and an accelerated wound healing rate, highlighting SPARC as a relevant factor in enhancing tissue regeneration within the model studied.

## DISCUSSION

The present study demonstrates that SPARC modulation in Wharton’s Jelly Mesenchymal Stem Cells (WJ-MSCs) profoundly influences their functional properties, secretome composition, and capacity to promote tissue regeneration. Through a combination of *in vitro* and *in vivo* assays, we provide evidence of SPARC’s pivotal role in regulating WJ-MSC secretory behavior, and their effects on fibroblasts, keratinocytes, and wound healing.

Our findings confirmed that SPARC expression in WJ-MSCs can be effectively modulated via lentiviral overexpression or knockdown without compromising their viability, morphology, or mesenchymal identity. Consistent with previous reports that SPARC regulates proliferation in other cell types^15–17^, SPARC knockdown in WJ-MSCs resulted in a significant increase in the proportion of cells in the G1 phase, suggesting a reduction in cell proliferation. Interestingly, SPARC overexpression did not significantly alter the cell cycle dynamics, potentially reflecting a context-specific role of SPARC in proliferation control.

Despite SPARC’s known role in cell adhesion and migration through its regulation of extracellular matrix (ECM) remodeling^11,18^, we observed no significant changes in the expression of key integrins on WJ-MSCs following SPARC modulation. This suggests that SPARC may influence WJ-MSC function primarily through secretome-related pathways rather than direct effects on adhesion molecules.

The secretome, consisting of soluble factors and extracellular vesicles, is a critical mediator of MSC-driven therapeutic effects^2,19,20^. Our results reveal that SPARC expression influences the biological activity of the WJ-MSC secretome. Conditioned medium (CM) from SPARC-modulated WJ-MSCs produced differential effects on keratinocyte and fibroblast behavior *in vitro*.

Keratinocytes exposed to CM from SPARC knockdown cells (KD-SPARC-CM) exhibited slower migration and wound closure compared to CM from wild-type (WT-CM) or SPARC-overexpressing cells (+SPARC-CM). This aligns with SPARC’s known role in promoting ECM remodeling and cell migration^21^. Fibroblasts, in contrast, displayed a higher proliferation rate when treated with KD-SPARC-CM, a response that may be tied to SPARC’s role in ECM composition and the modulation of growth factors such as transforming growth factor-beta (TGF-β)^22–24^. Interestingly, the effects of SPARC modulation were cell type-specific, highlighting the complexity of its role in tissue repair and regeneration.

Transcriptomic analysis revealed that SPARC modulates distinct gene networks in MSCs. SPARC knockdown induced a broader range of transcriptomic changes compared to overexpression, with several differentially expressed genes (DEGs) implicated in angiogenesis, inflammation, and extracellular matrix remodeling—hallmarks of tissue repair^25^. These findings are consistent with SPARC’s established roles in regulating ECM dynamics and tissue regeneration^11^. Notably, WNT2, a critical regulator of tissue regeneration^26^, was upregulated by SPARC and downregulated upon its knockdown, highlighting a potential key pathway by which SPARC influences regenerative functions. GO analysis further supported the role of SPARC in regulating genes involved in cell adhesion, response to growth factors, and angiogenesis.

The *in vivo* wound healing experiments demonstrated that CM from SPARC-overexpressing WJ-MSCs (+SPARC-CM) significantly accelerated wound closure and promoted tissue regeneration in mice compared to WT-CM and KD-SPARC-CM. Wounds treated with +SPARC-CM exhibited advanced epithelialization, reduced fibrotic tissue formation, and histological features resembling normal skin, including hair follicle regeneration. Conversely, KD-SPARC-CM treatment resulted in delayed healing, characterized by increased fibrotic tissue and incomplete remodeling of the dermis. These findings are consistent with SPARC’s known role in modulating collagen deposition and ECM remodeling and also with previous reports of impaired wound repair upon SPARC depletion^15,27,28^.

Interestingly, while the macroscopic and histological differences between treatment groups diminished by day 21, the quality of tissue regeneration varied. Wounds treated with +SPARC-CM exhibited superior ECM organization and reduced scarring, suggesting that SPARC plays a critical role in enhancing the quality of wound healing, in addition to its effects on healing speed.

Our results highlight SPARC as a key regulator of WJ-MSC function, particularly through its effects on the secretome. The enhanced regenerative properties observed with +SPARC-CM suggest that modulating SPARC expression could improve the therapeutic potential of MSC-based therapies for wound healing and tissue regeneration. Conversely, the delayed and fibrotic healing observed with KD-SPARC-CM underscores the importance of maintaining SPARC expression to prevent suboptimal outcomes.

Future studies should aim to identify the specific components of the SPARC-modulated secretome responsible for these effects, such as cytokines, growth factors, or extracellular vesicles. Additionally, investigating the role of SPARC in other MSC types or disease models could provide broader insights into its therapeutic potential.

## CONCLUSIONS

This study demonstrates that SPARC is a critical regulator of the regenerative functions of WJ-MSCs, primarily by modulating their secretome. SPARC overexpression enhances keratinocyte migration, promotes in vivo wound closure, and supports organized tissue regeneration, including hair follicle neogenesis. In contrast, SPARC knockdown impairs these functions and is associated with delayed healing, excessive fibrosis, and disorganized tissue repair.

These findings position SPARC not only as a marker of regeneration but as a functional enhancer of MSC-mediated wound healing. Future strategies that harness or mimic SPARC’s effects within MSC-based therapies may lead to more effective and targeted interventions for chronic wounds and other regenerative challenges.

## MATERIALS AND METHODS

### Human Umbilical Cord Obtainment and ethics

Written informed consent was obtained from each mother before normal cesarean birth. The donation is anonymous and the human umbilical cords were obtained from discarded placentas. All experiments and methods were performed in accordance with relevant guidelines and regulations. All experimental protocols and informed consents were approved by FLENI Ethics Committee.

### Isolation and Expansion of Wharton’s jelly Mesenchymal Stem Cells (WJ-MSCs)

For cell isolation, umbilical cords were transported in Dulbecco’s Modified Eagle Medium (DMEM, #11965092, Gibco). Each cord was chopped in 5 mm length, a sagittal cut was performed to expose Wharton’s jelly and umbilical blood vessels were carefully removed with clamps. Fragments were washed 2 or 3 times with Dulbecco’s phosphate-buffered saline (DPBS, Sigma-Aldrich) to remove the remaining blood. Then, the face with the exposed jelly was placed against the bottom of the culture plate and minimum essential medium (MEM-α, #11900073, Gibco), and 10% of fetal bovine serum (Natocor, Argentina). Plates were incubated at 37 °C in a humid atmosphere and 5% of carbon dioxide. Cell culture medium was changed every 2 to 3 days. WJ-MSC expansion was observed 10 to 14 days after explantation, and those cells were amplified until passage 2/3.

### Cell lines

Two types of human normal skin cell lines were used, human skin keratinocytes (HaCaT) and human skin fibroblasts (CCD-32Sk). Both cell lines were purchased from the Cell Culture Facility at UC Berkeley, California. CCD-32Sk cells were maintained in DMEM (#11965092, Gibco) supplemented with fetal bovine serum (Natocor, Argentina). HaCaT cells were cultured with DMEM supplemented with fetal bovine serum, sodium pyruvate (1mM, #11360070, Gibco) and MEM NEAA (#11140050, Gibco). Both cell lines were maintained at 37°C and 5% CO2.

### Plasmid Cloning of pLJM1-CMV-SPARC-Puro for SPARC Overexpression

To achieve SPARC overexpression, we generated the plasmid pLJM1-CMV-SPARC-Puro. This construct was based on the pLJM1-CMV-GFP plasmid (#19319, Addgene), which contains the necessary sequences for lentiviral packaging.

The SPARC mRNA sequence was amplified from an adenoviral plasmid, AAV2/2-CMV-SPARC, previously cloned in our laboratory. This adenoviral plasmid contained SPARC mRNA amplified from total RNA of CMM-WJ cells. We used Gibson assembly (New England Biolabs, NEB, US) for molecular cloning. The desired fragment was amplified by PCR using the following primers, designed according to the Gibson Assembly kit guidelines:

- Sparc assembly Forward: 5’-TGAACCGTCAGATCCGCTAGCCACTGAGGGTTCCCAGCAC-3’
- Sparc assembly Reverse: 5’-TGCCATTTGTCTCGAGGTCGAGAATTCGGCAAAGCTACAAATGGCAAGAG-3’

The amplified fragment was verified for size by gel electrophoresis and subsequently purified. The assembly reaction was performed by combining the digested pLJM1-CMV plasmid with the purified SPARC fragment at a 1:5 molar ratio. Gibson Assembly 2X Master Mix was then added, bringing the total reaction volume to 12 µL. The reaction was incubated in a thermocycler at 50°C for 30 minutes. Following the assembly, the entire reaction volume was used to transform competent Stbl3 bacteria.

**Plasmid Cloning of pLKO.1-shSPARC-Puro for SPARC Knockdown**

To achieve knockdown of SPARC, a short hairpin RNA (shRNA) targeting SPARC mRNA was designed using the GPP Web Portal, an online platform specialized in the design of gene-editing and gene-silencing sequences. The platform provides candidate shRNA sequences with an associated specificity score; sequences with a score above 7 are considered optimal. A specific shRNA sequence targeting SPARC, with a score of 15, was selected for cloning into the pLKO.1-Puro vector (#8453, Addgene). The following oligonucleotide sequences were used:

- shSPARC Forward: 5′- CCGGGAATACATTAACGGTGCTAAACTCGAGTTTAGCACCGTTAATGTATTCTTT TTG-3′
- shSPARC Reverse: 5′- AATTCAAAAAGAATACATTAACGGTGCTAAACTCGAGTTTAGCACCGTTAATGTA TTC-3′

For annealing, 1 µL of each oligonucleotide (100 µM) was mixed with 5 µL NEBuffer 2 (NEB) and Milli-Q water to 35 µL, incubated at 95°C for 3 minutes, and gradually cooled at 0.1°C/second to 16°C. In parallel, the pLKO.1-Puro plasmid was digested with AgeI-HF and EcoRI-HF (NEB) at 37°C for 1 hour, followed by gel purification.

Ligation was performed by mixing 20 ng of digested plasmid with 2 µL annealed oligos, 10 µL 2X Quick Ligase Buffer, and 1 µL Quick Ligase (NEB) in a 20 µL reaction, incubated at 16°C for 4 hours.

Then, 5 µL of the ligation mix was used to transform 50 µL of Stbl3 competent E. coli cells.

### Bacterial Transformation and Plasmid Verification

Competent E. coli Stbl3 cells (50 µL) were transformed with 1–3 µL of plasmid DNA, incubated on ice for 30 minutes, heat-shocked at 42°C for 90 seconds, and then placed on ice for 5 minutes. After adding 1 mL of LB medium, cells were incubated at 37°C for 30–60 minutes, centrifuged, resuspended, and plated on LB-agar plates containing the appropriate antibiotic. Plates were incubated overnight at 37°C.

Colonies were picked and grown in 3 mL LB + antibiotic at 37°C, 180 rpm for 16–18 hours. Plasmids were purified using the GeneJet Miniprep Kit (Thermo Fisher) and verified by sequencing (UC Berkeley DNA Sequencing Facility) and analyzed using Geneious software. Verified clones were expanded in 50 mL cultures, and plasmids were purified using the ZymoPURE II Midiprep Kit (Zymo Research).

### Lentiviral Particle Production and Transduction of WJ-MSCs

Lentiviral particles were generated using HEK 293T cells seeded at 3 × 10⁵ cells/well in 6-well plates with DMEM + 10% FBS. After 24 hours, at ∼60% confluency, the medium was replaced with serum-free DMEM, and cells were transfected using X-tremeGENE 9 (Roche) at a 3:1 reagent:DNA ratio. Transfection mixes were prepared by combining 9 μL of transfection reagent in 100 μL OptiMEM (Gibco) with 3 μg total plasmid DNA (1.5 μg gene-of-interest plasmid, 1.34 μg psPAX2, and 0.16 μg pMD2.G), incubated for 15–20 minutes, and added dropwise. After 24 hours, 10% FBS was added, followed by 1 mL DMEM + 10% FBS on day 3. Viral supernatants were collected on day 4, filtered (0.45 µm), and stored at −80°C.

For transduction, WJ-MSCs were seeded at 10,000–12,000 cells/cm². After 24 hours, medium was replaced with serum-free MEM-α, and 100 μL of viral supernatant was added dropwise. After another 24 hours, cells were cultured in MEM-α with 10% FBS and selected with 2 μg/mL puromycin. Medium was changed every 48 hours. Selection continued until all untransduced control cells were eliminated (48–72 hours). Transduced cells were maintained in standard culture medium for 2 additional days to ensure viral clearance.

### Membrane Protein Marker Analysis by Flow Cytometry

WJ-MSCs were detached and dissociated into single cells using TryPLE 1X (Gibco), diluted in medium, and centrifuged at 300 × g for 5 minutes. Cells were washed twice with PBS containing 0.1% BSA and stained in 50 µL of PBS/0.1% BSA with fluorophore-conjugated primary antibodies (1:50 dilution) at room temperature for 30 minutes.

The antibodies used were all from BD Pharmingen and included: CD90-FITC (Cat# 555595), CD73-PE (Cat# 550257), CD105-APC (Cat# 562408), CD49a-PE (Cat# 559596), CD49c-PE (Cat# 556025), CD49d-PE (Cat# 560972), CD49e-PE (Cat# 555617), CD49f-PE (Cat# 561894), CD14-FITC (Cat# 555397), CD34-FITC (Cat# 555821), CD45-FITC (Cat# 555482), HLA-DR-FITC (Cat# 555560), CD29-PE (Cat# 561795), and CD51/61-PE (Cat# 550037).

After staining, cells were washed twice and resuspended in 250 µL of PBS/0.1% BSA, then analyzed on a BD Accuri C6 flow cytometer. Unstained cells were used as controls for gating and to determine marker-positive populations. Data was analyzed using FlowJo software.

### Immunostaining and Fluorescence Microscopy

WJ-MSCs were cultured in 24-well plates and fixed with 4% paraformaldehyde in PBS for 20 minutes at room temperature. After fixation, cells were washed three times with PBS containing 0.1% BSA (Gibco). Permeabilization and blocking were performed using PBS with 0.1% Triton X-100 (Sigma), 0.1% BSA, and 10% normal goat serum (Gibco) for 45 minutes at room temperature.

Cells were incubated overnight at 4°C with anti-SPARC primary antibody (1:100 dilution; mouse monoclonal, Cat# 33-5500, Thermo Fisher) diluted in PBS with 0.1% BSA and 10% goat serum. After washing, cells were incubated for 45 minutes with Alexa Fluor 594-conjugated secondary antibody (1:400; goat anti-mouse, Cat# A11032, Invitrogen), and nuclei were counterstained with DAPI (200 ng/mL, Sigma, US).

Following final washes in PBS/0.1% BSA, cells were maintained in PBS, and fluorescence images were captured using an EVOS XL Core microscope and processed with ImageJ.

### Protein Analysis by Western Blot

Total proteins were extracted from cells using RIPA buffer (Sigma, US) with protease inhibitors (Calbiochem, US). Lysates were incubated on ice for 10 minutes and centrifuged at 10,000 rpm for 5 minutes. Protein concentration was determined using the Pierce BCA assay kit (Thermo Fisher, US), with BSA as standard.

Samples containing 30 µg of protein were denatured at 100°C for 5 minutes, separated by SDS-PAGE on 10% polyacrylamide gels (2.5 hours, 100 Volts (V)), and transferred to PVDF membranes (GE Life Sciences) for 45 minutes at 10 V. Membranes were blocked in PBS-Tween with 5% non-fat-dry milk for 1 hour and incubated overnight at 4°C with primary antibodies: mouse anti-SPARC (1:1000, Cat# 33-5500, Invitrogen) and rabbit anti-vinculin (1:1000, Cat# 129002, Abcam), diluted in PBS-Tween with 3% non-fat-dry milk.

After brief washes in PBS-Tween, membranes were incubated for 1 hour with HRP-conjugated secondary antibodies (Goat anti-rabbit and Goat anti-mouse, 1:5000, Dako), followed by chemiluminescent detection using SuperSignal West Femto (Thermo Fisher, US). Images were acquired with the ImageQuant LAS 4000mini system.

### Cell Cycle Analysis by Propidium Iodide (PI) Staining

To assess cell cycle distribution, DNA content was measured using propidium iodide (PI) staining. Cells were cultured to approximately 70% confluency, enzymatically dissociated with TryPLE, and resuspended in 1.5 mL of PBS. Fixation was performed by slowly adding 3.5 mL of cold 100% ethanol dropwise while gently vortexing, yielding a final ethanol concentration of 70%. Cells were fixed for 40 minutes at 4°C or stored at −20°C for later analysis.

After fixation, cells were centrifuged and washed three times with 1 mL of staining buffer (PBS + 3% FBS). The final pellet was resuspended in 500 µL of staining buffer, and 20 µL of RNase A (20 mg/mL, Thermo Fisher) was added. Samples were incubated for 30 minutes at 37°C. Then, 10 µL of PI solution (1 mg/mL, Sigma) was added, and cells were incubated for 5 minutes before acquisition.

DNA content was analyzed by flow cytometry using a BD Accuri C6 (Becton Dickinson). Cell cycle phases were quantified using FlowJo software with Watson model fitting.

### Conditioning of WJ-MSC Medium

WJ-MSC wild-type, +SPARC, and KD-SPARC cells were plated at a density of 10,000 cells/cm² in 60 mm culture dishes. After 24 hours, the culture medium was replaced with 4 mL of fresh MEM-α supplemented with 10% FBS (used for *in vivo* assays) or with DMEM supplemented with 10% FBS, sodium pyruvate (1mM) and MEM NEAA (for *in vitro* assays).

Following 48 hours of incubation for conditioning, the medium was harvested and centrifuged at 4000 rpm for 15 minutes at 4°C.

### *In Vitro* Wound Healing Assay

HaCaT cells (180,000 cells per well) were seeded into 24-well plates and allowed to adhere for 16–18 hours before initiating the assay. The next day, a scratch was made through the confluent cell monolayer using a 10 µL pipette tip. After two PBS washes to remove debris, 1 mL of conditioned medium (CM) was added to each well. The treatment groups included CM from WJ-MSC wild-type (WT-CM), WJ-MSC overexpressing SPARC (+SPARC-CM), and WJ-MSC with SPARC knockdown (KD-SPARC-CM). A control group received the standard medium used for conditioning WJ-MSCs (DMEM supplemented with 10% FBS, sodium pyruvate (1mM) and MEM NEAA.

Plates were placed in a BioTek Lionheart FX automated microscope (Agilent) and maintained at 37°C with 5% CO₂. Time-lapse imaging was performed using Gen5 software (version 3.11), capturing images every 30 minutes for 14 hours at 4X magnification in both brightfield and phase contrast modes.

Image analysis was conducted to assess wound closure over time by quantifying the cell-free area. Percent wound closure was calculated for each condition, and statistical comparisons were made to evaluate differences between treatment groups.

### Animal model and ethics

Immunocompetent adult mice of the C3H/S strain, inbred in the Biotherium of the Histology Chair of the Faculty of Medical Sciences, were used. This strain has been maintained by inbreeding in the aforementioned facility since 1966, following the arrival of the original breeders sent by Professor J. W. Wilson (Department of Biology, Brown University, Providence, Rhode Island, USA). Mice were housed in individual boxes under a 12-hour light/12-hour dark cycle (Circadian Rhythms Room), with forced ventilation, controlled temperature (22 ± 2 °C), and ad libitum access to food and water.

All animal procedures were conducted in collaboration with Dr. Marcela N. García from the Cytology, Histology, and Embryology Department, Faculty of Medical Sciences, Universidad Nacional de La Plata (UNLP). This study was approved by the UNLP ethics committee (COBIMED, Protocol: P01-03-2018). All procedures were conducted in accordance with institutional and international guidelines for animal care and use. Anesthesia was induced using a ketamine-diazepam combination administered intraperitoneally, and animals were monitored for signs of adequate sedation. Euthanasia was performed via an overdose of anesthetic agents followed by a secondary method to ensure death, in compliance with AVMA Guidelines for the Euthanasia of Animals (2020 edition).

This study complies with the ARRIVE guidelines (Animal Research: Reporting of In Vivo Experiments) 2.0 to ensure rigorous and transparent reporting of animal research.

### *In vivo* Wound Healing Assay

Immunocompetent C3HS mice (four animals per condition, two males and two females) were used for all experiments, which were performed in triplicate. Mice were anesthetized with a combination of Ketamine (100 mg/kg, Vet Pharma) and Diazepam (5 mg/kg, Lamar) and skin wounds were executed with a disposable dermatological biopsy punch. Incisions were made through the epidermis, dermis, and subcutaneous tissue, leaving the deep fascia intact. A week later, a secondary intentional wound was made at the same site, designated as day 0. Immediately following the secondary wounding, the first dose of conditioned medium (CM) was administered. Three types of CM were used: wild type (WT-CM), +SPARC-CM (derived from WJ-MSC overexpressing SPARC), or KD-SPARC-CM (derived from WJ-MSC with knockdown of SPARC). Each animal received a daily intradermal injection of 200 µL of CM, containing 100 µg of protein, for 10 consecutive days. For the first experiment, an additional control group was included and treated with unconditioned medium (MEM-α supplemented with 10% FBS) to assess whether the presence of serum in the medium had any significant effect on wound healing. As the effect was notably greater in the mice treated with conditioned media derived from mesenchymal cells, the subsequent two experiments were carried out using only the groups treated with conditioned media, in order to minimize the use of experimental animals.

Daily observation of the wound was performed by 2 independent observers. Photographs of the wounds were taken on day 0 (day of secondary wounding) and on days 7, 10, and 21 post-injuries. Wound areas were subsequently analyzed using ImageJ (FIJI) to calculate wound closure rates. At each experimental time point, animals were deeply anesthetized via intraperitoneal injection of ketamine (100 mg/kg) and diazepam (5 mg/kg). Upon confirmation of a surgical plane of anesthesia—evidenced by the absence of pedal and corneal reflexes—euthanasia was performed by decapitation using sterile instruments. Wound tissue samples were immediately collected and processed for histological analysis. Sections were stained with hematoxylin and eosin (H&E) to assess general tissue structure and cellularity. Then, Masson’s trichrome staining was used to determine the degree of connective tissue regeneration, especially collagen fibers. All histological analyses were performed by specialized technicians within the collaborative group led by Dr. García.

### Whole Transcriptome Sequencing (RNA-seq)

Total RNA was extracted using TRIzol reagent (Invitrogen) following the manufacturer’s instructions and quantified with Qubit Fluorometric Quantitation (Thermo Fisher). 5 µg of RNA per sample were treated with DNaseI to remove genomic DNA contamination and subsequently precipitated with 0.1 volumes of 3 M sodium acetate (pH 5.5) and 2 volumes of 100% ethanol. RNA quality and integrity were verified by automated electrophoresis before sequencing. Library preparation and RNA sequencing were carried out by Macrogen (South Korea) using the TruSeq Stranded Total RNA with Ribo-Zero kit (Illumina).

Raw sequencing reads (FASTQ files) were aligned to the human reference genome (GRCh38/hg38) using the STAR aligner (v2.7). Gene-level count quantification was performed with HTSeq-count, and downstream differential gene expression analysis was conducted using the DESeq2 package in R (v4.1.2). Genes exhibiting a fold-change greater than 1.5 or smaller than -1.5 and an adjusted p-value (FDR) < 0.05 were considered significantly differentially expressed.

To visualize the global transcriptional differences across conditions, a hierarchical clustering heatmap was generated using the Seaborn package in Python (3.10). Differentially expressed genes across all comparisons were standardized using Z-score transformation of normalized gene expression values. Clustering was performed based on Euclidean distance and complete linkage to reveal patterns of gene expression similarity among samples and conditions.

Gene Ontology (GO) enrichment analysis was performed using ShinyGO (v0.76.3) with default parameters. The list of significantly differentially expressed genes was used as input to identify enriched biological processes, molecular functions, and cellular components. Enrichment was calculated using the hypergeometric test with FDR correction, and significantly overrepresented GO terms (adjusted p-value < 0.05) were reported. Pathway enrichment analyses based on KEGG databases were also performed to identify significantly enriched signaling and metabolic pathways.

### Statistical Analysis

Statistical significance of the observed differences was analyzed using GraphPad PRISM version 8. A Student’s t-test with Welch’s correction was employed to compare differences between indicated treatments. Differences were considered statistically significant when p-values were less than 0.05 (p < 0.05).

## Acknowledgements

We would like to thank Fleni Institute-CONICET and Pérez Companc Foundation for their continuous support.

## Author contributions statement

AL carried out data collection, data analysis and writing of the original draft, JS, contribute with data collection, analysis and interpretation of RNA-seq data, writing and critical revision of the article, MBP, AI and MNG, contributed with data collection, analysis and interpretation of the *in vivo* experiments. CL devised the project and design the study and assisted with drafting and critical revision of the manuscript. GS, SM and CL conceived the original idea and helped supervise the project. All authors discussed the results and contributed to the final manuscript.

## Data Availability

The data discussed in this publication have been deposited in NCBI’s Gene Expression Omnibus (Edgar et al., 2002) and are accessible through GEO Series accession number GSE301307 (https://www.ncbi.nlm.nih.gov/geo/query/acc.cgi?acc=GSE301307).

## Conflict of Interest Statement

The authors declare that they have no competing interests.

## Notes

### Competing Interest Statement

The authors have declared no competing interest.

