## Supplementary data for "Modulating SPARC Expression in Mesenchymal Stem Cells Improves Secretome-Mediated Skin Regeneration and Wound Repair"

a

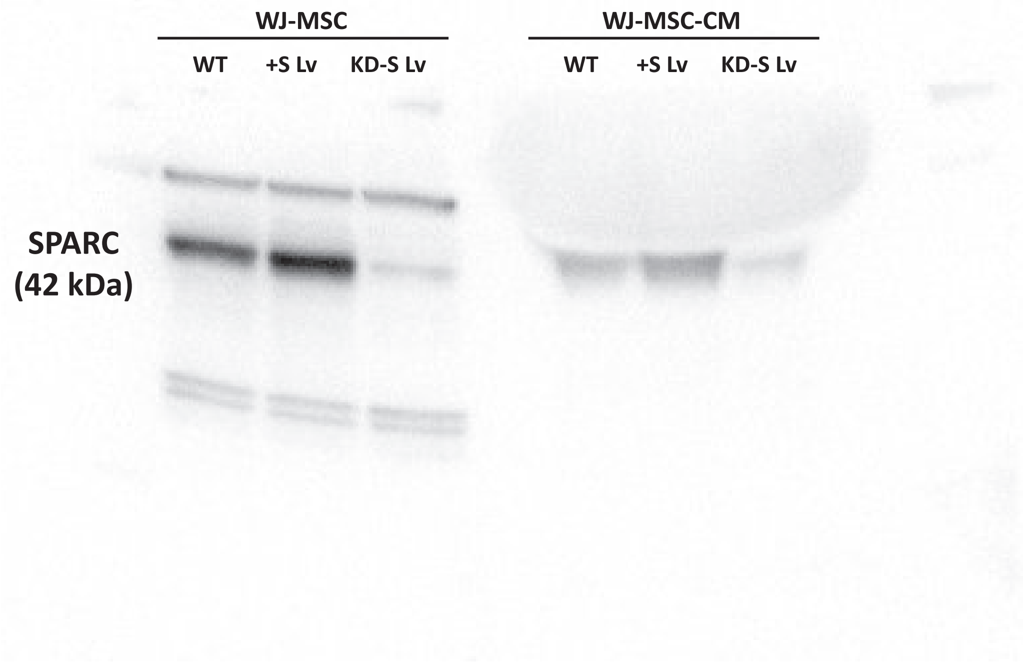
WJ-MSC
WJ-MSC-CM
WT
+S Lv
KD-S Lv
WT
+S Lv
KD-S Lv
SPARC
(42 kDa)

b

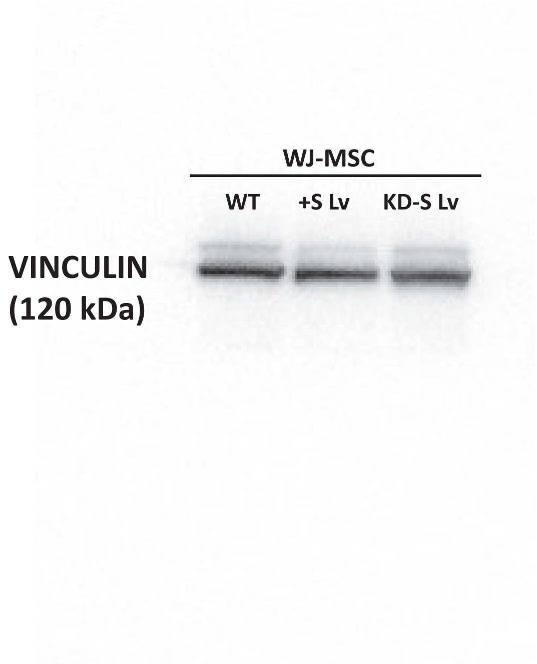
WJ-MSC
WT
+S Lv
KD-S Lv
VINCULIN
(120 kDa)

**Supplementary Figure 1. SPARC expression levels.**

**a:** SPARC expression levels in WJ-MSC lysates (left panel and graphic) and their conditioned media (CM) (right panel and graphic) assayed by SDS-PAGE and Western blot. Cell lysates and CM were obtained from wild-type (WT) WJ-MSCs or those transduced with +SPARC Lv or KD-SPARC Lv. Whole blot is shown. **b:** VINCULIN expression levels, used as housekeeping protein, in WJ-MSC lysates assayed by SDS-PAGE and Western blot. Cell lysates were obtained from wild-type (WT) WJ-MSCs or those transduced with +SPARC Lv or KD-SPARC Lv. Whole blot is shown.

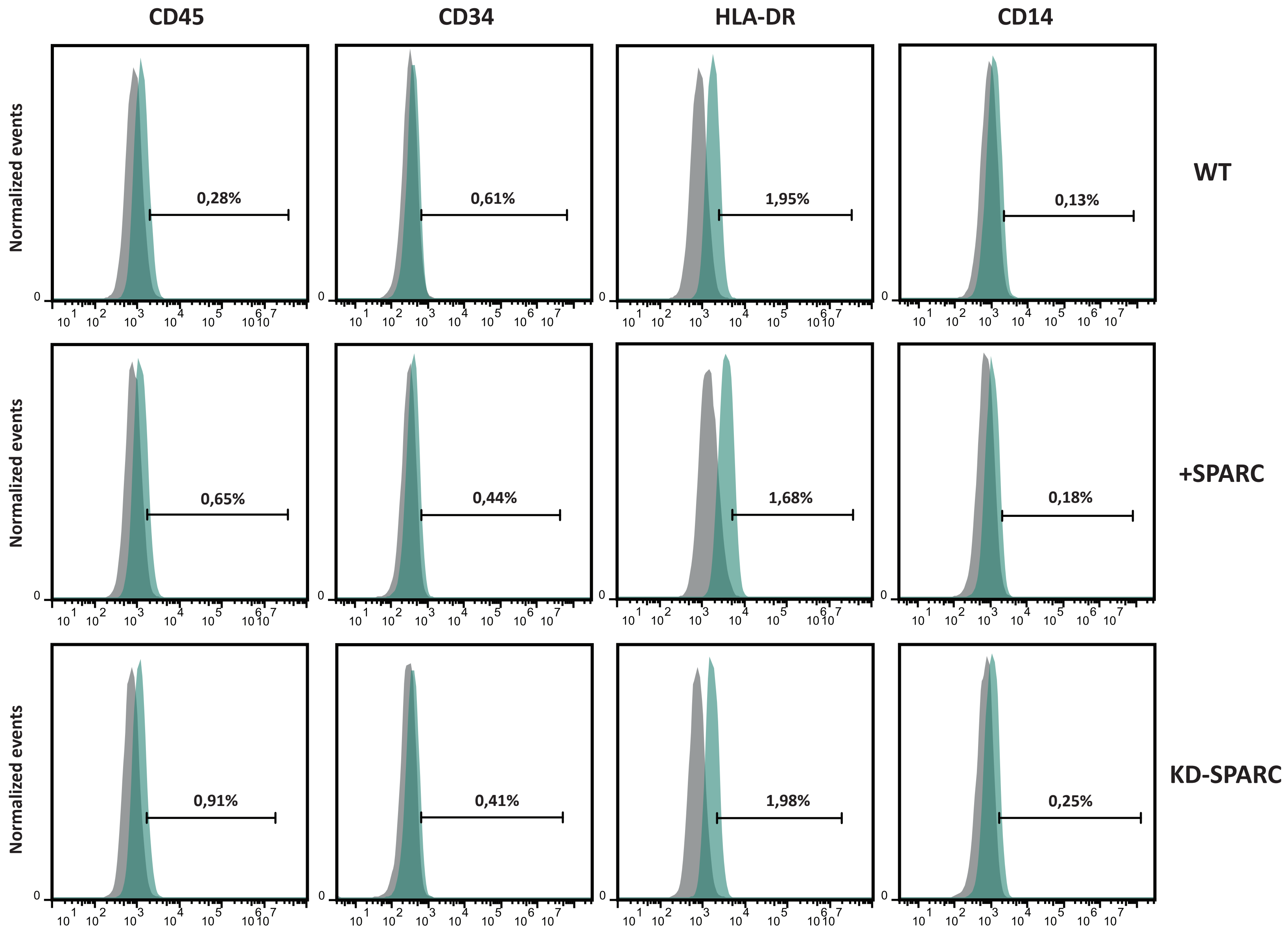
CD45
CD34
HLA-DR
CD14
WT
+SPARC
KD-SPARC
Normalized events
0,28%
0,61%
1,95%
0,13%
0,65%
0,44%
1,68%
0,18%
0,91%
0,41%
1,98%
0,25%

**Supplementary Figure 2. WJ-MSC characterization.**

MSC negative markers CD45, CD34, HLA-DR and CD14, measured by flow cytometry. Representative plots of three independent experiments. Plots are representative of three biological replicates. Percentages were calculated with the FlowJo v10.0 software. Statistical analysis was made with Graphpad 8.0 software.

**Supplementary video 1.** Wound healing assay of HaCaT cells treated with UM for 14 hours.

**Supplementary video 2.** Wound healing assay of HaCaT cells treated with WT-CM for 14 hours.

**Supplementary video 3.** Wound healing assay of HaCaT cells treated with +SPARC-CM for 14 hours.

**Supplementary video 4.** Wound healing assay of HaCaT cells treated with KD-SPARC-CM for 14 hours.

**Table 1.** Differential expressed genes in +SPARC *vs*. WT condition.

| Gene name | Log2 Fold Change | P value |
|---|---|---|
| SORCS1 | 4,088836547 | 0,000261204 |
| ELN | 2,824752453 | 0,007107983 |
| GABRB1 | 1,850383057 | 0,04913346 |
| CMKLR1 | 1,525709093 | 0,0000134 |
| SOCS2 | 1,276314093 | 0,0000134 |
| GFRA1 | 1,050226186 | 0,00000525 |
| GALNT3 | 1,042839968 | 0,00606799 |
| FENDRR | 0,915204784 | 0,004906428 |
| WNT2 | 0,858442942 | 0,040589884 |
| DUSP5 | 0,857458893 | 0,008567076 |
| ELAPOR2 | 0,825468566 | 0,013848857 |
| CCBE1 | 0,771724674 | 0,001785832 |
| FAT3 | 0,732592668 | 0,010265648 |
| NLGN1 | 0,677922358 | 0,040589884 |
| IGFBP5 | 0,642728725 | 0,000219594 |
| CHRM2 | 0,586531222 | 0,005789346 |
| GATA6 | -0,718246344 | 0,043739895 |
| CPXM1 | -0,753913254 | 0,001981101 |
| EGR1 | -0,840067647 | 0,042480843 |
| NYNRIN | -0,981343825 | 0,003901288 |
| EDA2R | -0,988889953 | 0,040589884 |
| FOS | -1,071627998 | 2,87E-11 |
| GAS7 | -1,0831385 | 0,003901288 |
| PRUNE2 | -1,105652583 | 0,001204953 |
| FOSB | -1,107168466 | 2,09E-08 |
| ABCA8 | -1,236415872 | 0,0000409 |
| SULF2 | -1,467211329 | 0,010265648 |
| PTGIS | -1,487900069 | 0,011411777 |
| UNC5B | -1,667515441 | 0,004070398 |

**Table 2.** Differential expressed genes in KD-SPARC *vs*. WT condition.

| Gene name | Log2 FoldChange | P value |
|---|---|---|
| MCF2L2 | 5,076736503 | 0,024030597 |
| TPD52 | 4,80878534 | 0,004213229 |
| GAD1 | 4,477008071 | 0,023760141 |
| ADGRL3 | 4,372201947 | 0,025093924 |
| BEX1 | 3,526555304 | 2,61E-17 |
| CA3 | 3,006121369 | 0,021110637 |
| MDGA2 | 2,871778102 | 0,032965243 |
| EPDR1 | 2,806018474 | 0,02903465 |
| EPB41L3 | 2,578009602 | 0,012966764 |
| DGAT2 | 2,547741588 | 0,008400497 |
| ANKRD1 | 2,439823667 | 0,000194828 |
| LCP1 | 2,435096701 | 8,02E-11 |
| AK4 | 2,352145807 | 0,028368524 |
| PGM5 | 2,313511021 | 0,016424287 |
| ITGA7 | 2,238462839 | 0,012056346 |
| CCNA1 | 2,206443034 | 0,023317435 |
| LRP1B | 2,071352841 | 0,023474636 |
| JAKMIP2 | 2,051572852 | 0,0000768 |
| ABLIM1 | 2,000114866 | 0,000003 |
| PRKAA2 | 1,929376182 | 0,018995897 |
| DYSF | 1,800084028 | 0,036871186 |
| ATF3 | 1,768863266 | 0,000105298 |
| MAMDC2 | 1,62977303 | 0,010880934 |
| PTPRB | 1,620531983 | 0,000186146 |
| DCLK1 | 1,579821807 | 0,023760141 |
| NCAM1 | 1,503903289 | 0,000000211 |
| EPHA3 | 1,402409164 | 0,030426271 |
| PTPN3 | 1,377715875 | 0,001181291 |
| GALNT3 | 1,356461497 | 0,0000475 |
| TINAGL1 | 1,354053803 | 0,004619869 |
| KRT7 | 1,3454117 | 0,038056087 |
| FAT3 | 1,343195751 | 3,84E-11 |
| TNFRSF21 | 1,324755234 | 0,022243202 |
| CHST11 | 1,321114364 | 0,012266768 |
| PLCE1 | 1,289041199 | 0,010972897 |
| HES1 | 1,285324023 | 0,004619869 |
| TLL1 | 1,278839207 | 0,04995493 |
| WWC1 | 1,230259936 | 0,024030597 |
| NRK | 1,230224966 | 0,037933611 |
| TAPBPL | 1,220723734 | 0,025455593 |
| DDIT4L | 1,18718454 | 0,012630709 |
| SH3RF2 | 1,177089665 | 0,002434818 |
| PTPRH | 1,163587661 | 0,038056087 |

| | | |
|---|---|---|
| LYPD6B | 1,158125287 | 0,003417094 |
| BIRC3 | 1,137881015 | 0,0000516 |
| IL1A | 1,111373177 | 0,004212496 |
| PLAAT3 | 1,058103247 | 0,023474636 |
| CXCL8 | 1,039071071 | 0,007468448 |
| DUSP1 | 1,033405237 | 0,000000139 |
| CARD11 | 1,006673878 | 0,000768027 |
| IRAK3 | 0,978542783 | 0,000431764 |
| ALDH1A3 | 0,970254384 | 0,000000117 |
| ARHGDIB | 0,942920062 | 0,000335736 |
| CYFIP2 | 0,925266218 | 0,006678846 |
| MFSD6 | 0,897142141 | 0,001585113 |
| MACROH2A2 | 0,860915235 | 0,012056346 |
| MYEF2 | 0,851681279 | 0,004619869 |
| ZC3H12C | 0,812753776 | 0,008539867 |
| IL1B | 0,806418739 | 0,000625627 |
| SERPINE1 | 0,796798187 | 0,0000149 |
| JAG1 | 0,793205013 | 0,011876453 |
| FAR2 | 0,789453607 | 0,044623481 |
| C3 | 0,787256053 | 0,013192016 |
| SERPINB7 | 0,776747529 | 0,004922606 |
| NABP1 | 0,775001278 | 0,0000149 |
| LYPD6 | 0,760925614 | 0,013797326 |
| HDAC9 | 0,750692597 | 0,010026326 |
| PDCD1LG2 | 0,734476634 | 0,017363055 |
| TNFAIP3 | 0,734454209 | 0,00344609 |
| PIGV | 0,731398059 | 0,038056087 |
| HBEGF | 0,730398443 | 0,004619869 |
| ABLIM3 | 0,703891712 | 0,034120038 |
| EPHB2 | 0,676135259 | 0,006200776 |
| TGFB2 | 0,658193622 | 0,018537019 |
| C3orf52 | 0,649621655 | 0,017601228 |
| RCAN1 | 0,617721629 | 0,008128602 |
| ABL2 | 0,587126056 | 0,019917141 |
| CHN1 | -0,596428208 | 0,040166825 |
| HSPA2 | -0,599977862 | 0,005351755 |
| FMNL2 | -0,630409904 | 0,002130446 |
| EPS8 | -0,650200521 | 0,000768027 |
| THBS2 | -0,662280299 | 0,022981529 |
| C1R | -0,667023725 | 0,005198635 |
| TBX2 | -0,669833921 | 0,027982268 |
| PLOD2 | -0,700691157 | 0,00002 |
| PDGFRA | -0,720126141 | 0,003435998 |
| ACVR2A | -0,723720325 | 0,044623481 |

| | | |
|---|---|---|
| PRICKLE1 | -0,733369911 | 0,045889538 |
| CPT1A | -0,740518137 | 0,016949027 |
| PCDH9 | -0,748239192 | 0,022908223 |
| WFDC21P | -0,765161631 | 0,00516468 |
| LTBP1 | -0,765243775 | 0,028069843 |
| NRP1 | -0,766601441 | 0,00000389 |
| SLC25A30 | -0,785822743 | 0,040945634 |
| PDZRN3 | -0,814769979 | 0,003110162 |
| AP1AR | -0,819569638 | 0,024030597 |
| TSHZ1 | -0,833464528 | 0,035752683 |
| TBCC | -0,839825986 | 0,00516468 |
| LMNB1 | -0,867291117 | 0,000778659 |
| ALDH1L2 | -0,87322803 | 0,018720182 |
| PPM1F | -0,890156342 | 0,00000594 |
| SLC14A1 | -0,96060293 | 0,00387378 |
| NPTN | -0,9732756 | 1,28E-08 |
| TRIB2 | -0,973479264 | 0,038056087 |
| DLG3 | -1,002035298 | 0,00387378 |
| CNTNAP3B | -1,026106312 | 0,00387378 |
| FSCN1 | -1,06547316 | 2,67E-13 |
| FBLN5 | -1,089957108 | 0,011034956 |
| PCDH18 | -1,12049598 | 0,002927663 |
| PDE7B | -1,125094512 | 0,000268795 |
| GAS1 | -1,157445049 | 0,0000516 |
| HGF | -1,188422665 | 0,008539867 |
| PTGIS | -1,224458255 | 0,016949027 |
| PTGER2 | -1,334518917 | 0,001413325 |
| ALDH1A1 | -1,388869448 | 0,000830489 |
| DCN | -1,511616333 | 6,13E-10 |
| DDIT4 | -1,523244915 | 0,002533906 |
| WNT2 | -1,557907644 | 0,001193079 |
| BMP6 | -1,612002791 | 0,002533906 |
| AQP1 | -1,723658177 | 0,00387378 |
| TMEM119 | -2,374484392 | 6,13E-10 |
| SPARC | -2,55661936 | 8,21E-90 |
| TDO2 | -2,588665921 | 0,037148165 |
